# *Leishmania* Ribosomal Protein (RP) paralogous genes compensate each other’s expression maintaining protein native levels

**DOI:** 10.1101/2023.09.15.557908

**Authors:** Francisca S. Borges, José C. Quilles, Lucas B. Lorenzon, Caroline R. Espada, Felipe Freitas-Castro, Tânia P. A. Defina, Fabíola B. Holetz, Angela K. Cruz

**Author notes:** These authors contributed equally as first authors (Francisca S. Borges, José C. Quilles).

## Abstract

In the protozoan parasite Leishmania, most of the genes encoding for ribosomal proteins (RPs) are present as two or more copies in the genome, their untranslated regions (UTRs) are predominantly divergent, and might be associated with a distinct regulation of the paralogous genes’ expression. Here, we investigated the expression profiles of two RPs (S16 and L13a) encoded by duplicated genes in Leishmania major. The genes encoding for S16 protein have identical CDSs and divergent UTRs while the L13a CDSs diverge in two amino acids with divergent UTRs. Using CRISPR/Cas9 genomic editing system, we generated knockout (Δ) and tagged transfectants for each paralog of L13a and S16. Combining tagged and Δ cell lines we show that the expression of both RPS16 and RPL13a isoforms differ throughout the parasite development with one of the isoforms being always more abundant than its respective copy. Additionally, compensatory expression was observed for each paralog when one of the isoforms was deleted, evidencing functional conservation of these proteins. Such phenomenon is related to post-translational processes, since the compensation happened at the protein levels, with no alterations observed at the transcript levels. Ribosomal profiles for RPL13a point out a standard behavior for these paralogues as already reported for other RPs in trypanosomatids, showing its interaction with heavy RNA-protein complexes. The identification of sets of proteins binding specifically to the 3’UTRs of either the high or less abundant transcripts suggests a possible role of these proteins to differently control the levels of expression of these RP genes. In addition, conserved cis-elements were identified in the 3’UTRs of RPS16 or RPL13a; among these, exclusive cis-elements for the more or for the less expressed transcripts were identified.

## INTRODUCTION

Ribosomal proteins (RPs) are essential ribosome constituents in all organisms. In eukaryotes, 46 and 33 different RPs are present in the ribosome large (RLS) and small (RSS) subunits, respectively (1). In some organisms, such as yeast and fungi, most genes encoding for RPs are duplicated as identical or very similar copies (2). Genome duplication and hybridization, retro-duplication and polyploidy have been the most discussed concepts used to explain the emergence of the duplication phenomenon (3). Comparing duplicated genes, identical or very similar protein coding DNA sequences (CDSs) with divergent untranslated regions (*UTR*s) suggests a distinct and individual regulation of gene expression (4), while dissimilarities in the CDSs may be associated with functional differences among the paralogs (5). However, information on the expression control of these genes in different organisms is still scarce.

Besides their orthodox function, noncanonical activities of RPs have been reported in different cellular pathways and organisms (6–8), such as disturbing the bud site selection (9) and cell development in yeast (10). More recently, RPs were shown to be also involved in signaling pathways and development of cancer cells (11). In colorectal cancer, for instance, the inactivation or partial mutation of the isoform RPL22 promoted the upregulation of its homologous paralog RPL22L1, leading to increased drug resistance (12). Furthermore, noncanonical functions have been attributed to RPL13a, a conserved eukaryotic RP that regulates the translation of specific mRNAs molecules in humans (13).

Ribosomes are highly conserved RNA-protein complexes that are responsible for translation (14). Differences in the levels of ribosomal subunits (RS) may directly affect translation rates in the cell, and alterations in the expression of paralog RP genes directly affects RS levels. In yeast, the two paralogues of both *RPL26* and *RPL33* genes share a highly conserved CDSs (> 98%), but in both cases only one of the two paralogs exerted a positive influence on RLS abundance (10). Studies in other organisms reporting different functions for duplicated genes encoding ribosomal proteins, demonstration of moonlight activities for RPs and the fact that most of the RP genes are duplicated in the genome of *Leishmania* led us to investigate if two duplicated RP genes, both with divergent *UTR*s and one with two amino acid substitutions, play different roles in *L. major*.

In trypanosomatids all protein coding genes are organized in polycistronic units (PTUs) (15) lacking canonical promoters for the individual gene transcription (16). As a consequence, the control of gene expression in these organisms occurs mainly at the posttranscriptional level (17), depending on mRNA transport, stability, decay and rate of translation (18). *Leishmania* parasites are adaptive organisms switching between mammalian and invertebrate hosts during their life cycle, and modulating gene expression for their adaptation and survival in the distinct hostile environments (19). Co-transcriptionally, mRNAs are trans-spliced at 5’*UTR* in a process coupled with the polyadenylation of the upstream gene (20). Gene expression modulation in these parasites is strongly dependent on *cis* elements present at the 3’*UTR*, which are recognized by RNA binding proteins (RBPs) that direct the transcript to different fates (21). Interestingly, the translation of RP genes in trypanosomatids has been recently reported as regulated by different proteins binding to their 5’ or 3’*UTR*s (22). Thus, divergences in the *UTR*s of duplicated genes may be key for their differential gene expression regulation at the level of transcript stability and/or protein translation control. In *Leishmania*, multicopy genes are frequent (15), and the players involved in their differential expression are largely unknown. To shed light on this regulation, we here studied profiles of expression of two duplicated RP genes with divergences only in the *UTR* (*RPS16* genes) or both, in the *UTR* and CDS (*RPL13a* genes).

## MATERIALS AND METHODS

### Sequence Alignment

*L. braziliensis* RPL13 and RPS16 protein and gene sequences were obtained from TriTryp data base (https://tritrypdb.org/tritrypdb/app/) and applied as multiple global alignments queries in Clustal Omega (https://www.ebi.ac.uk/Tools/msa/clustalo/) to obtain the final comparison.

### Parasite culture, differentiation, and transfection

Procyclic promastigotes of *Leishmania major* strain LV39 (MRHO/SU/59/P) were cultivated in M199 medium (Sigma Aldrich) supplemented with 10% heat-inactivated fetal bovine serum. To obtain metacyclic promastigotes, procyclic forms were cultivated for 5 days in M199 and metacyclics were enriched from stationary phase cultures using a Ficoll^®^ gradient (23). Transfections were performed using the AMAXA Nucleofector equipment, as already published by our group (24). Briefly, 1·10^7^ promastigotes were resuspended in 100 µL Tb-BSF buffer and added to 60 µL of DNA generated by PCR (for Δ or tagging). All this volume was transferred to an electrolytic cuvette and transfection was performed using the X-001 program of Amaxa Nucleofector instrument (LONZA). The culture was maintained in M199 at 26°C before selection with the appropriate drugs (16μg/mL hygromycin-B; 20μg/mL blasticidin; 16μg/ mL puromycin) on solid M199 media. After 15-20 days, individual colonies were collected and transferred to liquid M199 medium containing the respective selection drug and homozygosity was confirmed by DNA extraction and conventional PCR.

### Immunofluorescence

The subcellular location of RPs was analyzed by immunofluorescence assays: a total of 1.5×10^6^ cells was centrifuged at 1,400 x g for 5 minutes at RT, followed by washing with 500 μl of PBS. Cells were fixed for 10 minutes at RT in 500 µL of 3% paraformaldehyde in PBS, pelleted and washed once with 500 μL PBS. The pellet was resuspended in 100 μL of 0.1% glycine in PBS and 30 μL of the total cells were added to poly-lysine slides and left to adhere for 30 minutes.

Then, fixed cells were permeabilized with 30 μl of 0.2% TRITON in PBS solution for 5 minutes at RT, followed by five times PBS wash steps. Blocking was performed for 30 min with 5% skim powder milk in TBS-T. Anti-*Myc* (Sigma C3956) primary antibody was diluted in the same blocking solution at 1:4,000 ratio and incubated for two hours. After that, five washes were performed with PBS and secondary antibodies conjugated with Alexa Fluor 488 (Invitrogen A11001) were 1:500 diluted in PBS and incubated for 30 minutes at RT protected from the light. Nuclei and kinetoplast staining were performed using HOESCHT (Invitrogen H3570) at 2 µg·mL^-1^ for 15 min. Images were acquired on a Leica DMI6000B fluorescence microscope at 60x magnification and processed using the Fiji Image software J (https://imagej.net/Fiji/Downloads).

### Scanning Electron Microscopy (SEM)

10^7^ parasites were fixed for 2 hours in 3% paraformaldehyde and 2% glutaraldehyde in PBS supplemented with 0.9 mM CaCl_2_ and 0.5 mM MgCl_2_ at RT. The parasites were post-fixed in 2% OsO_4_ for 2 h and incubated with a thiocarbohydrazide (TCH) solution for 10 minutes, followed by ethanol and acetone dehydration. Then, parasite cells were mounted on a support and subjected to gold-coated metal plating. Cells were analyzed using an electron microscope scan (JEOL-JSM-5200). Images were captured by an Electron Microscopy Multiuser Laboratory (Department of Cellular and Molecular Biology, Faculty of Medicine of Ribeirão Preto, USP).

### Western Blotting

Parasites were pelleted (1,400 x g for 5 minutes at RT), washed once with 500 µL of cold-PBS supplemented with protease inhibitors (Roche), resuspended in 10 µL of extraction buffer^54^ and boiled for 10 min. Total protein was quantified in Nanodrop One spectrophotometer (Thermo Scientific) by measuring the absorbance at 280 nm. For every protein sample, sample buffer was added (24) and boiled for 3 minutes. 30 ug of total protein were fractionated in a 12% polyacrylamide gel. Proteins were transferred to nitrocellulose membranes (GE Healthcare Life Sciences: 10600003) and blocked for 1 h with TBS-T buffer (Tris-saline-Tween buffer: 10 mM Tris, 100 mM NaCl, pH 7.6, Tween20 0.1%) containing 5% milk powder. Immune detection was performed with the appropriate primary and secondary antibodies, following manufacturer’s recommendations: α-Myc (1:4,000, Sigma C3956) and α -EF1α: (1:80,000, Merck 05-235). Both primary antibodies were diluted in solution of TBS-T with 2.5% powdered milk and incubated for 1 h at RT, followed by incubation with secondary antibody at same conditions. Membrane visualization was performed by the chemiluminescent substrate interaction (ECL kit – GE Healthcare: RPN2232) and images were obtained on the ImageQuant LAS 4000 equipment (GE Healthcare). Band quantification was performed by Fiji ImageJ software comparing the band skew between the samples.

### RNA extraction and transcripts quantification

Cells were pelleted (1,400 x g for 5 minutes at RT), lysed with TRIzol reagent (Invitogen) and RNA extraction was performed using DirectZol RNA Miniprep kit (Zyme Research). Total RNA was treated with DNase Turbo (ThermoFisher Scientific) and RT-qPCR experiment was performed and analyzed as described by Freitas Castro and cols (25), using G6PDH as housekeeping genes for normalization.

### Pull-down assay

The regions corresponding to the 3’*UTR* of the *RPS16* and *RPL13a* genes were retrieved from TriTryp data base (https://tritrypdb.org/tritrypdb/app/). These sequences were cloned into pUC-56 plasmid between a T7 promoter and 4xS1m aptamer sequences, adapted from Leppek’s strategy (26). Then, RNA was *in vitro* transcribed (MEGAscript T7 transcription kit – ThermoFisher AM1334). Then, 30 µg of purified RNA was immobilized on Streptavidin magnetic beads (NEB) at 4°C during 8h under orbital rotation. 10^8^ parasites were lysed on ice by physical pressure using a 19G needle with 1mL of SA-RNP-Lyse buffer (20mM Tris-HCl pH7.5, 150mM NaCl, 1.5mM MgCl_2_, 2mM DTT, 2mM RNase inhibitor, 1 protease inhibitor cocktail tablet, 1% Triton X-100). Biotinylated proteins were previously removed from the extract by incubating the lysed extract for 8 h at 4°C with the streptavidin beads. Then, the supernatant was incubated to the RNA sequences immobilized on the magnetic beads during 8h at 4°C. An empty plasmid with no sequence between T7 promoter and 4xS1m aptamer was used as RNA control to identify the unspecific proteins. After that, beads were washed three times with wash buffer (20mM Tris-HCl pH7.5, 300mM NaCl, 5mM MgCl_2_, 2mM DTT, 2mM RNase inhibitor, 1 protease inhibitor cocktail tablet), resuspended in 35 µL of Laemmli buffer (27) and boiled for 10 min before application into 12% polyacrylamide gel. Samples were run at 110V until the samples reached the separation gel.

### Proteomic analysis, Mass Spectrometry, DataBase Searching and Criteria for protein identification

Gel bands containing the samples were sent to protein identification by mass spectrometry analysis (Proteomics Platform of the CHU de Quebéc Research Centre, Quebec, Canada). Three biological replicates were evaluated for each protein and for the control. Results were obtained and analyzed using the software Scaffold Protein. The list of identified proteins was filtered using a protein threshold of 99%, a peptide threshold of 95% and a minimum of 1 peptide identified for all the samples. Proteins interacting with the control RNA sequence were unconsidered and the results were based on the proteins specifically interacting with the 3’*UTR* sequences in triplicate. Detailed information on Mass Spectrometry, Database searching and criteria for protein identification have been described elsewhere (24).

### Starvation resistance assay

Nutritional stress was evaluated by incubating 10^6^ parasites per well in a 96 well plate in PBS for 4 h. After that, plate was centrifuged 1,400 x g for 5 minutes at RT, cells were resuspended in complete M199 with MTT 1mg·mL^-1^ and incubated for 24 h at 26°C. After that, parasites were pelleted and 200 µL of DMSO was added to solubilize the tetrazolium crystals. Absorbance was measured at 560 nm considering the substrate conversion for cell non-starved as 100% of nutritional response. Experiments were biologically performed in triplicate with quintuplicate technical replicates.

### Sucrose density gradient

Promastigotes polysomes from Leishmania major under normal and nutritional stress conditions were purified and fractionated on sucrose gradient (28). Briefly, 10^9^ cells were previously incubated or not in PBS 100% for 2 h and then cycloheximide 100 µg·mL^-1^ was added for 10 min. Cells were kept on ice and washed once with TKM buffer supplemented with 100 µg·mL^-1^ cycloheximide, 10 µg·mL^-1^ heparin, 10 µg·mL^-1^ E-64 and protease inhibitor cocktail (Roche). For polysome dissociation, cells were treated with puromycin 2mM. Cells were pelleted and 100 µL of buffer lysis (TKM supplemented with 10% surfactant NP- 40 and 2 M sucrose), lysed by up and down agitation followed by centrifugation at 18,000 x g at 4°C for 10 min. The supernatant was added on the top of linear 10-50% sucrose density gradient prepared in the same buffer. The system was centrifuged at 39,000 rpm at 4°C for 2 h in a Beckman SW41 rotor. After centrifugation, the gradient fractions were collected by ISCO gradient fractionation system, which 30 µL of each sample was used for blotting assays.

### Statistical Analysis

Statistical t-test (Mann–Whitney) and one-way ANOVA followed by Tukey’s multiple comparison tests were performed using GraphPad Prism 8, considering *p-*value <0.05 as statistically significance.

## RESULTS

### RPs from paralog genes have different expression levels and undistinguishable subcellular distribution

We investigated the expression of two pairs of duplicated paralogous genes encoding for ribosomal proteins in *L. major* LV39 strain: RPS16 (Fig1A) and RPL13a (Fig1B). *RPS16* was studied as a model for paralogs with identical coding sequences and divergent *UTR*s (Fig1C), while *RPL13a* is a model for paralogs with non-identical coding sequences and divergent *UTR*s (Fig1D). Seven nucleotide substitutions were found when comparing the two copies of *RPL13a* genes; two of them led to a codon modification and to a non-conserved amino acid substitution (Fig1D). Notwithstanding, we here focused on the role of the divergent 3’*UTR*s, which are known to be deeply involved in the interaction with RNA binding proteins (29) and, therefore, in the control of gene expression by regulating or modulating the mRNAs transcriptional and translational rates, as well as their stability.

**Figure 1.**
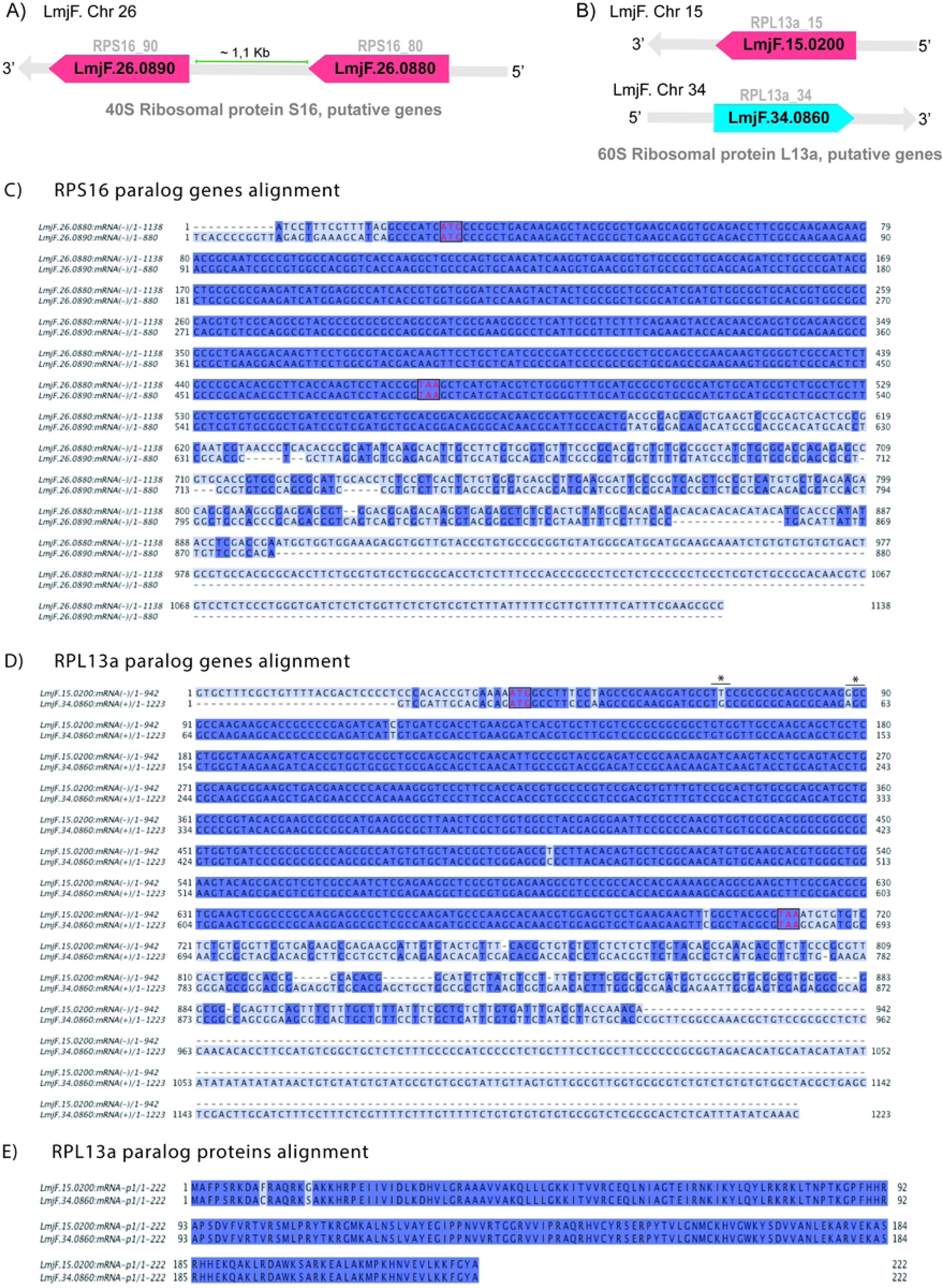
Paralog genes and proteins alignments. (A) 40S ribosomal protein S16 genes are located in tandem in the minus strand of chromosome 26 and (B) 60S ribosomal protein L13a genes are found in distinct chromosomes (15 and 34) and in opposite directions. (C) RPS16 and (D) RPL13a gene alignments, highlighting the divergences in the UTR sequences. (E) RPL13a amino acid sequence alignment showing the two divergent amino acids.

To investigate the fate of each of these RPs, we used CRISPR/Cas9 to tag each one of the proteins at N-terminus following a well-established protocol (30). A cassette containing 3x *myc* epitopes and the blasticidin resistance gene was used to repair the Cas9-induced double strand break at 5’-end of each one of the paralogs (Fig2A). The insertion of the tag in each copy of the RPS16 paralogs was confirmed by PCR, showing that each individual paralog, RPS16_80 or RPS16_90, was efficiently tagged in both alleles (FigS1). For the RPL13a isoforms, because of the similar amplicon sizes, the PCR was carried out for both copies, *RPL13_15* and *RPL13_34*, allowing the confirmation of tag insertion in only one of them, and with the maintenance of unmodified paralogous copy (FigS1). These transfectants were used to check the individual expression level and subcellular localization of all four RP isoforms.

**Figure 2.**
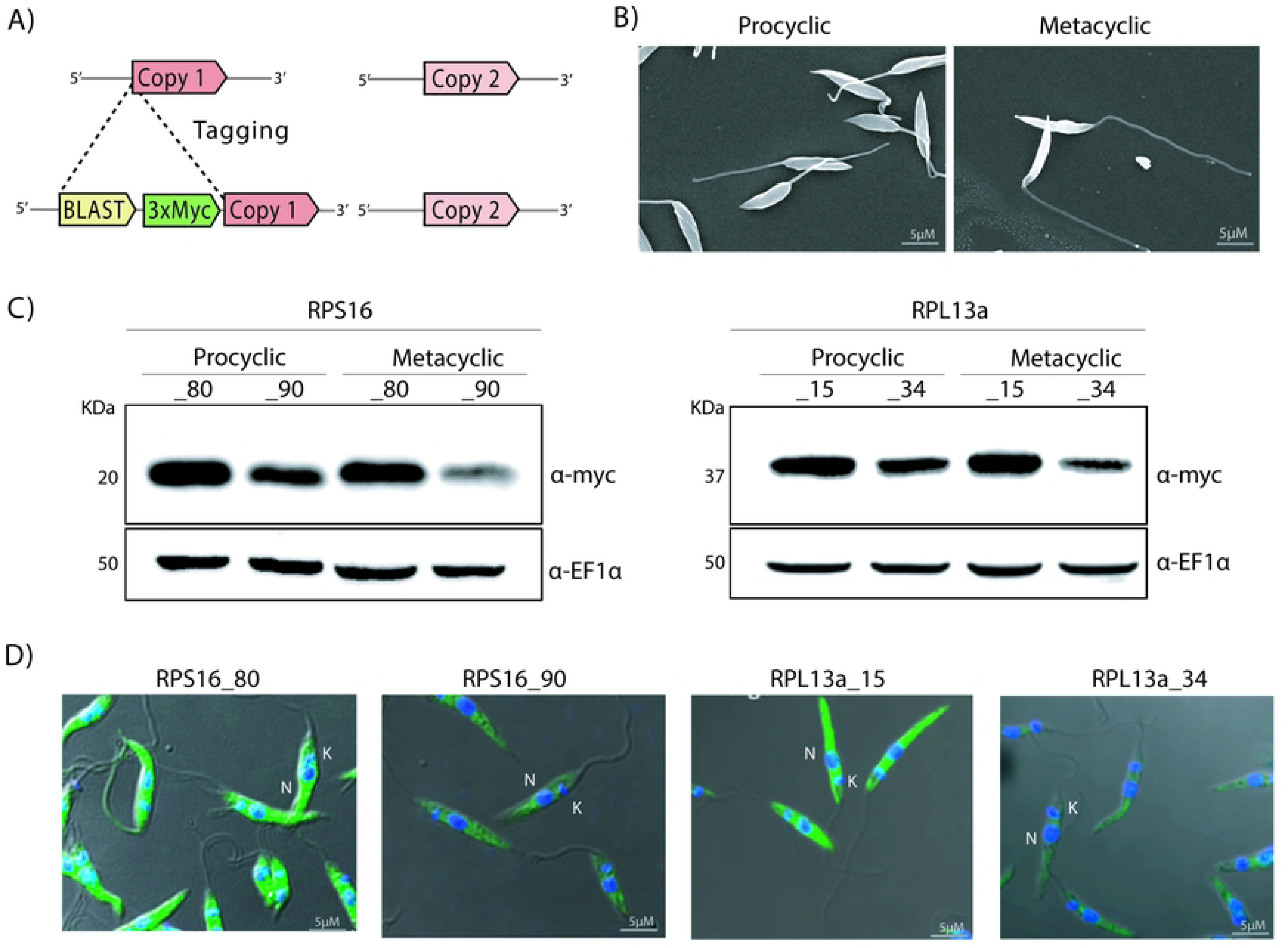
Parental protein levels of duplicated S16 and L13a ribosomal proteins. (A) Strategy for 3xmyc tag insertion in only one of the copies of the ribosomal protein genes using CRISPR/Cas9. (B) Procyclic promastigotes were cultivated until the log phase from which metacyclic forms were purified by Ficoll gradient (see methods), and their morphological differences were confirmed by scanning electron microscopically. (C) Western blotting using α-myc showed that, for both ribosomal proteins, one of the copies is always less expressed than the other, and is further less abundant in metacyclic compared to procyclic promastigotes. (D) Immunofluorescence confirmed the cytoplasmic localization for all RP (green) copies in procyclic promastigotes (nucleus (N) and kinetoplast (K) are indicated and stained in blue using Hoechst).

Log-phase procyclic and metacyclic promastigotes (purified from cultures in stationary phase by a Ficoll gradient fractionation (23) were isolated from axenic culture and their morphologies were confirmed by scanning electron microscopy considering differences in the cell body volume and flagellum size, since metacyclic promastigotes bear an elongated flagella and reduced cell body compared to procyclics (Fig2B).

The relative protein abundance derived from each paralogous gene was evaluated by western blotting (Fig2C) in procyclic and metacyclic promastigotes. RPS16_80 and RPL13_15 were consistently present in higher levels than their respective paralogous proteins in both morphologies. Such difference was more evident between RPS16_80 and RPS16_90 isoforms in metacyclic promastigotes; RPS16_80 levels are unchanged between procyclic and metacyclic stages and RPS16_90 is always lower than RPS16_80 but the difference between RPS16_80 e RPS16_90 levels is more marked in metacyclic forms.

Protein localization was investigated by immunofluorescence using an anti-*myc* antibody (Fig2D). As expected, the RPs were found exclusively in the cytoplasm, with no detectable co-localization with DAPI in the nucleus or in the kinetoplast. Also, a similar analysis for RPS16 subcellular distribution using an α-RPS16 antibody in the parental *L. major* LV39 confirmed the findings with the *myc*-tagged versions of RPS16 (FigS2). Importantly, the intensity of the immunofluorescence signal corroborated the western blotting results and confirmed the differences in the abundance of each RP isoform in procyclic promastigotes.

### Duplicated RPs present a paralog compensation that maintains protein levels

Next, we sought to check if different paralogs could compensate for the absence of each other in a compensatory mechanism that maintains protein at parental levels. To this end, cells carrying a *myc* tag in a given paralogous gene was knocked out for its respective paralog counterpart (Fig3A). All Δ were confirmed by PCR (FigSF3) and the protein levels were evaluated in both procyclic and metacyclic promastigotes by western blotting using a α-*myc* antibody. For all the studied proteins, when one of the genes was deleted, the tagged paralog protein levels increased and likely compensated for the lack of its counterpart (Fig3B). This compensatory effect is more evident when the gene that is expressed at lower levels (*RPS16_90* and *RPL13_34*) is kept and their highly expressed paralog is knocked out (Fig2C). This phenomenon was observed in both procyclic and metacyclic stages and for both pairs of genes. Interestingly, the maintenance of each ribosomal protein levels seems to be important even at the metacyclic stage, which presents lower translational and metabolic activity.

**Figure 3.**
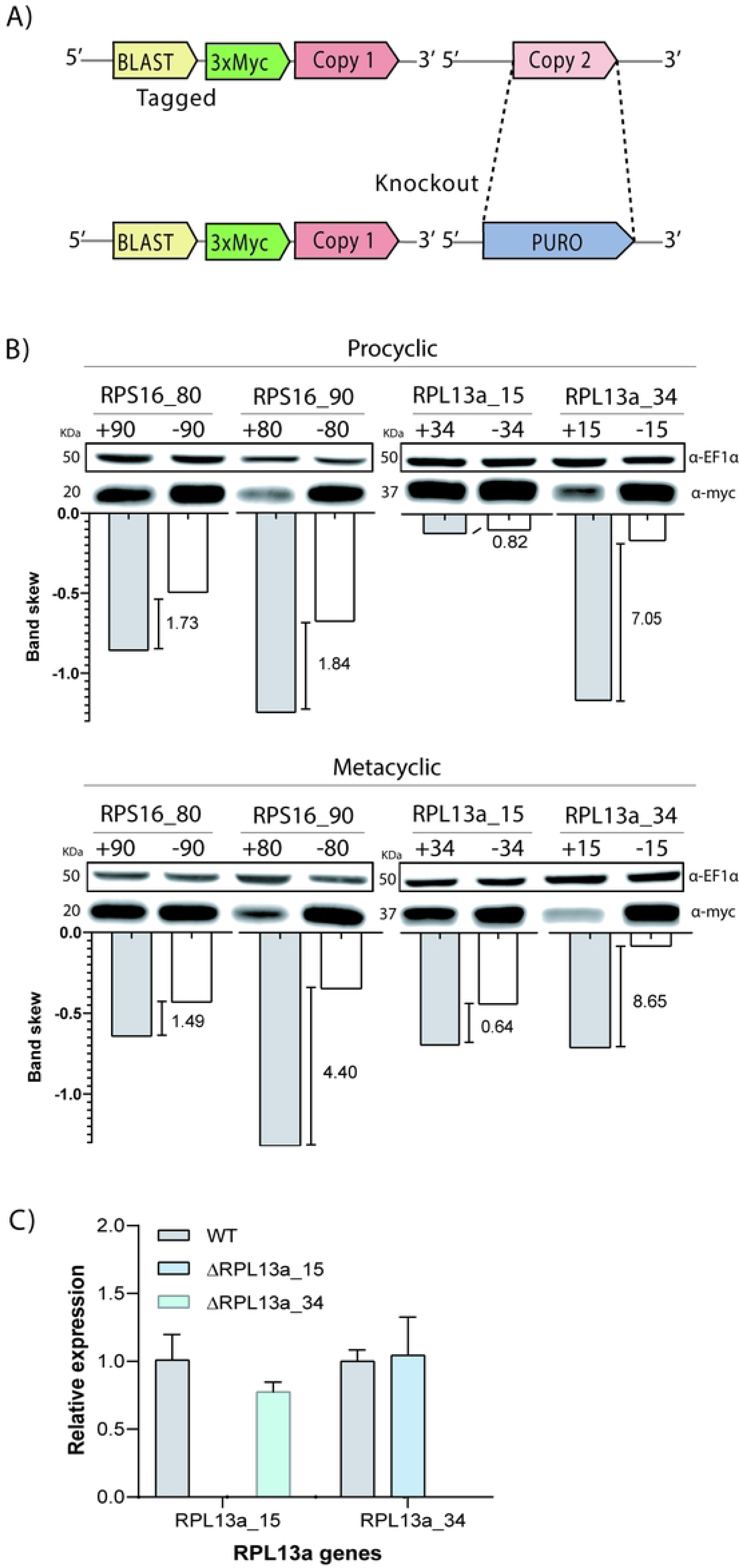
Compensatory expression of RP paralogs in knockout cells. (A) After tagging one of the RP copies, the other one was deleted using CRISPR/Cas9 and (B) the expression levels in procyclic and metacyclic promastigotes were analysed by western blotting using α-myc. The presence and absence (knockout) of the correspond RPS16 and RPL13a paralogues are indicated by + and -, respectively. Quantification of band intensity was calculated by imageJ and the results were plotted in the graphic with the increment of band colour indicated. (C) Quantitative PCR was used to quantify the relative expression of RPL13a genes in the parental (WT) and knockout (Δ) parasites and were calibrated to the G6PDH housekeeping gene.

Also, to confirm that the presence of at least one of them was necessary for parasite viability, we pursued knocking out both *RPS16* paralogs, but no double Δ clones were recovered: despite having inserted two drug resistance genes in two rounds of transfections using a CRISPR/Cas9 system, all transfectants retained at least one copy of a parental gene intact (FigS3).

In addition to protein amounts, gene expression was also evaluated at the transcript levels to check at what level(s) the compensatory mechanism occurs. For that, total RNA from *L13* Δ procyclic promastigotes was used in RT-qPCR utilizing specific primers for the divergent *UTR* regions of each L13 gene copy comparing transcript levels in the Δ and parental cell lines. Relative expression was normalized based on the expression of the housekeeping gene Glucose-6- phosphate dehydrogenase (G6PDH). No significant differences at transcript levels were observed for either *RPL13a* paralog (Fig3C). This result indicates that RPL13a compensatory mechanism involves either translation rate or protein stability, but it is not related to increases in transcript levels and stability.

### Profile of proteins interacting with the UTRs of the studied RP transcripts

Since the 3’*UTR* of both duplicated genes are not conserved and a compensatory expression mechanism was observed, we performed pull-down assays using S1m *in vitro* system (26) to identify potential proteins involved in such mechanism. Briefly, the 3’*UTR* sequences were determined based on TriTrypDB annotations (Fig4A) and fused to the S1m aptamer. After immobilization in streptavidin-coated magnetic beads, the 3’*UTR* sequences were incubated with log-phase *L. major* LV39 protein extract. The bound proteins were isolated and identified by mass spectrometry (MS). Gene ontology (GO) analysis revealed that the proteins interacting with *RPL13a* and *RPS16* 3’*UTR*s are mostly involved in peptide and ribosome biogenesis, respectively (Fig 4B). Among these proteins, 6 were identified as binding to all four 3‘*UTR* sequences (Fig4C), and four of them are directly related to protein folding and ribosomal biogenesis and processing. We may speculate that these four proteins that commonly bind to all the examined *UTR*s are possibly core proteins binding to mRNA of RP genes and might be involved in intra nuclear trafficking and nucleolar activities (Fig4D).

**Figure 4.**
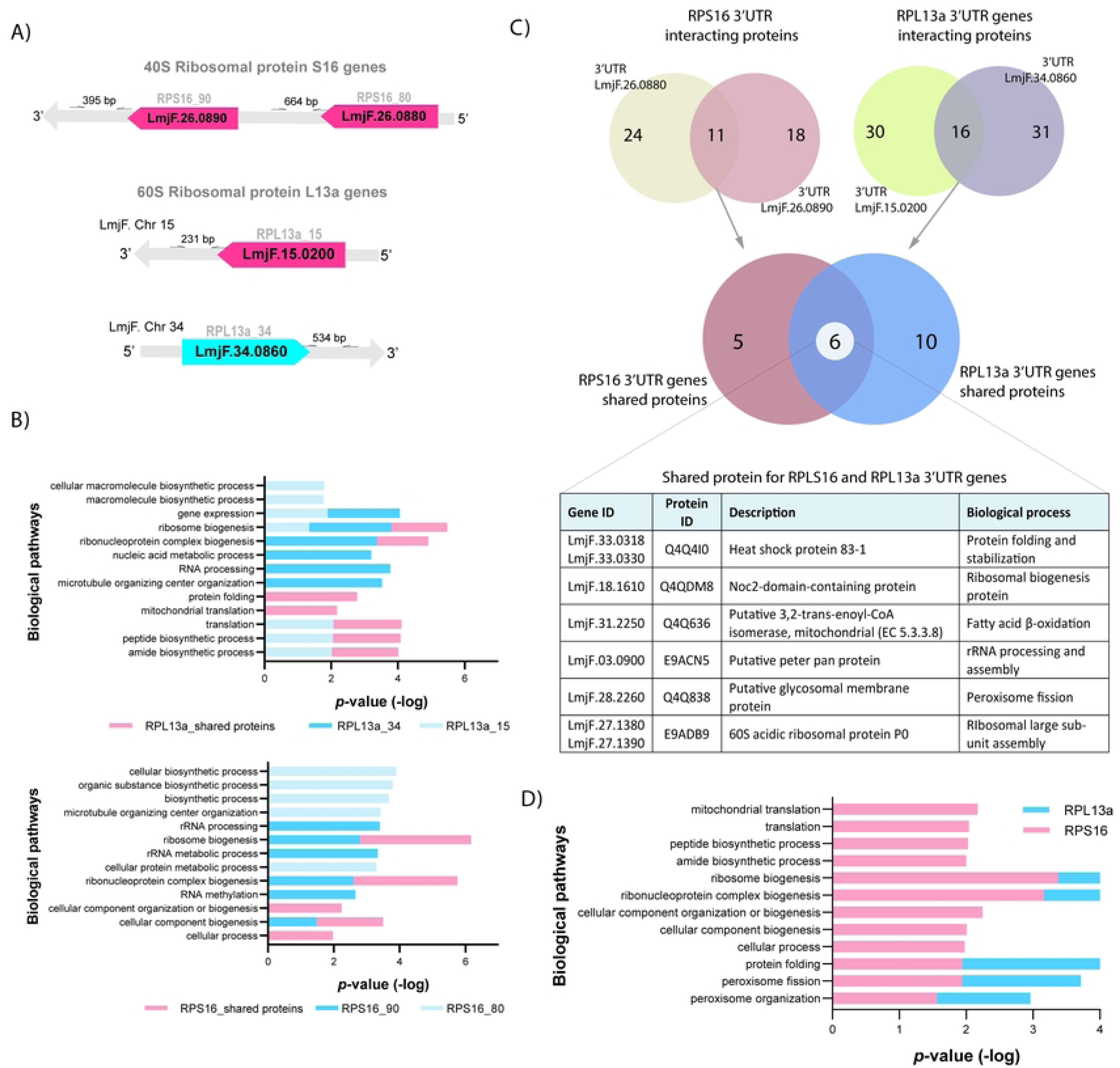
Identification of proteins interacting with the 3’UTRs of the RPS16 and RPL13a paralogs mRNA. (A) Based on the reads for each gene, the 3’UTR for each gene was amplified by PCR and cloned into PUC-54-4xs1m plasmid for pulldown assay. (B) Gene ontology analysis of proteins identified in vitro interacting with individual 3’UTR of each RPS16 and RPL13a paralog (shades of blue) genes, and proteins which interacted with both paralogs (shared - red). (C) Specific and shared proteins for each duplicated genes were identified, with total of 24 and 18 for RPS16_80 and 90, respectively. For RPL13a paralogues, similar numbers of specific proteins were obtained: 30 and 31 proteins for RPL13a_15 and 34, respectively. Altogether, six proteins were found binding to all four 3’UTRs of the duplicated genes encoding for RPS16 and RPL13a proteins; three of them are involved in ribosomal functions. (D) All 3’UTRs of ribosomal protein duplicated genes bound to proteins involved in ribosome pathways, when considered the five most relevant p-values (D). All the analyses were based on the results from three independent assays.

### Increased response to starvation and RP levels of mutant parasites

Considering that ribosomal activity is affected under nutritional stress potentiating or uncovering non-canonical functions of ribosomal proteins, as shown for other organisms (6–9), we analyzed the resistance to starvation comparing parental cells to RP transfectants knocked out for each one of the paralogs. The ability to rapidly adapt and respond to starvation is critical for *Leishmania* differentiation and survival. Procyclic promastigotes were incubated for 4h in PBS, with recovery of 24h in fresh medium, followed by colorimetric assays using MTT (Fig5A).

**Figure 5.**
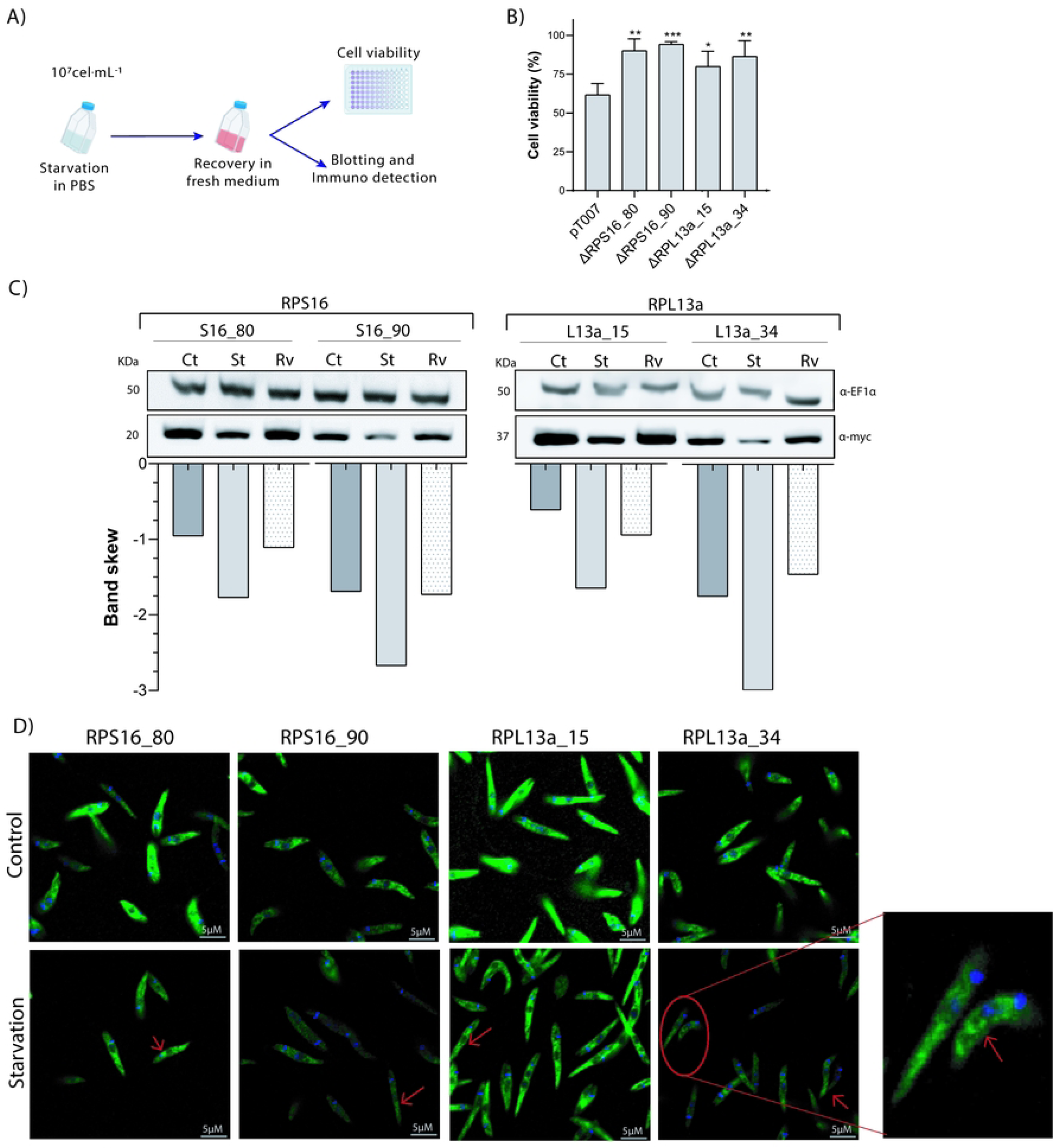
Nutritional stress response and expression levels of RPs under starvation. (A) Procyclic promastigotes in exponential phase of growth were exposed to total starvation in PBS for 4h at 27°C, followed by viability assay, blotting and immunodetection. (B) Cell viability was quantified by MTT assay at 24h post-starvation, checking the recovery capacity of the mutant cells compared to parental line. (C) Expression levels of RPs were lower under starvation (St), particularly for RPS16_90 and RPL13_34, with general recovery of parental levels after 24h of recovery in fresh medium (Rv) in comparison to the non-starved cells (control – Ct) – detection by western blotting with α-myc. (D) Immunofluorescence revealed no significant different between the RPs distribution (green stain) under stress, but a subtle accumulation can be observed by increment of signal in some regions of the cytoplasm (arrows) after 4h of total starvation (D).

Curiously, an increment of the parasite resistance to the nutritional stress environment was observed for all the Δ parasites compared to the parental line, as indicated by the higher mitochondrial activity measured by MTT conversion (Fig5B).

Additionally, the RP levels were evaluated by western blotting prior to and after 4 hours of starvation in PBS, as well as at 24hs of recovery in fresh medium. As shown (Fig5C), upon starvation, a subtle decrease in the levels of both L13 and S16 proteins was observed, which was completely recovered to original levels after 24h in fresh medium. Interestingly, this trend was also observed for RPS16_90 and RLP13_34 which are always expressed at lower levels under physiological conditions (Fig5C). Additionally, at stress conditions no marked changes in the subcellular distribution of these RPs were observed (Fig5D).

### RPL13a accumulates in non-polysomic fractions under starvation

Polysomic profile analyses are informative tools to investigate ribosomal subunits assembly status, translation activity and biomolecule complexes (28,31). Given that RPL13a proteins are not identical with two non-conserved amino acids substitutions at the N-terminus (Fig1E), we investigated the ribosome organization of these Δ cells under physiological and starvation conditions. Parental and Δ cells bearing only one of the *RPL13a* paralogs were kept under physiological or nutritional stress (2h) conditions and their cell extracts were fractionated by a sucrose gradient after treatment with cycloheximide or puromycin. For comparison purposes, we first analyzed the profile of polysomic distribution in parental (pT007) and Δ cells under normal and stress conditions in the presence of cycloheximide and observed a subtle decrease in the levels of the 80S complex for the Δ*RPL13a_15* under the non-starving condition (Fig6A). However, after 2 hours in PBS, both Δ*RPL13a_15* and Δ*RPL13a_34* parasites presented lower pre-polysome content compared to the parental cell line. Next, we analyzed the distribution of each RPL13a paralog throughout the non-polysomic and polysomic fractions. Under physiological conditions, there was no clear distinction in the distribution of a specific paralog in the absence of the other, with both paralog proteins being detected at similar levels at sucrose densities corresponding to monosomes and light or heavy polysomes (Fig6B). Although no polysomes were detected after starvation in the cell extracts analyzed, both RPL13a paralogs were detected mainly at the density of light polysome fractions when treated with puromycin, as if large and small subunits were assembled but not organized for translation (Fig6C).

**Figure 6.**
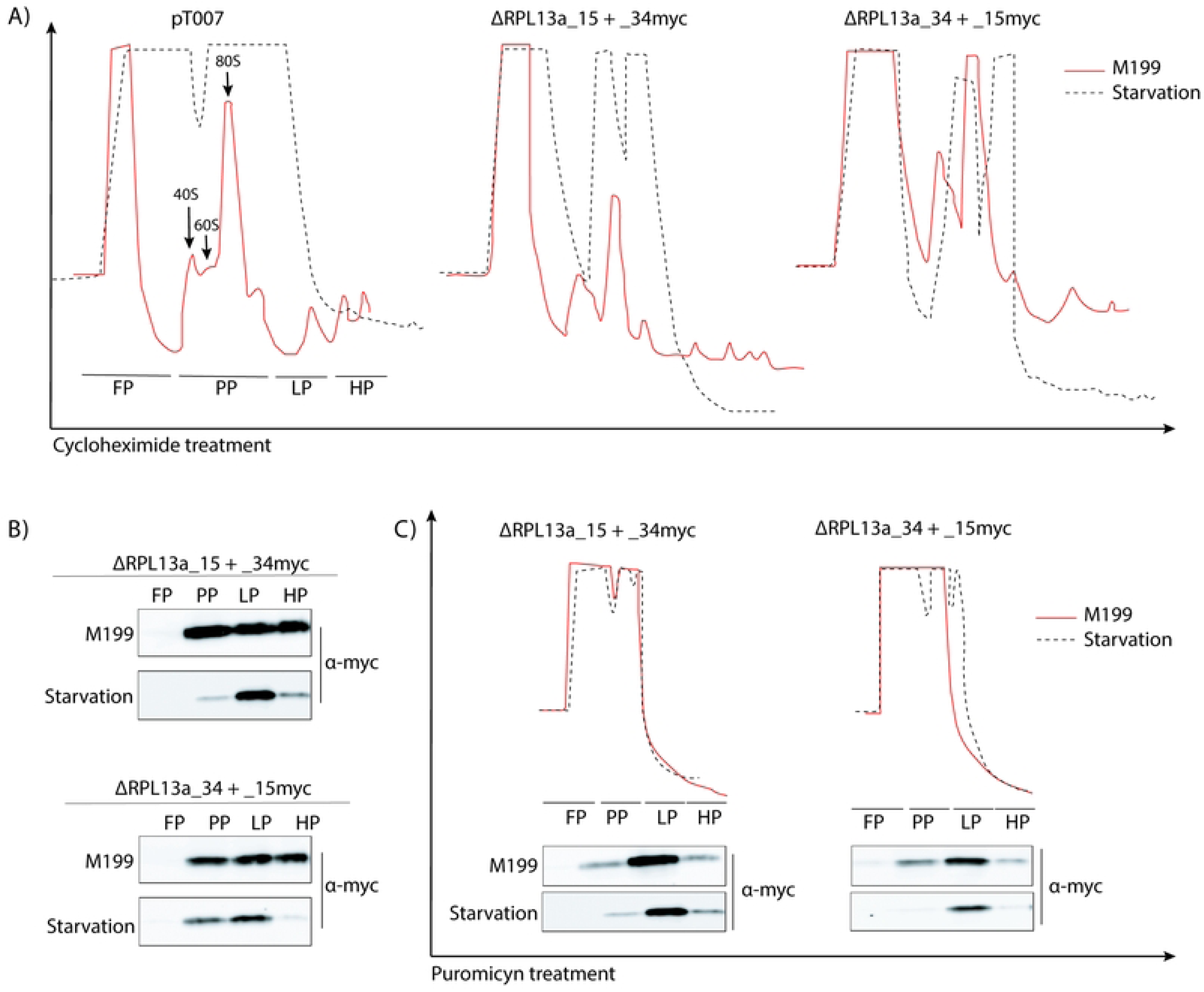
Alteration in the polysome profile by nutritional stress. (A) RPL13a Δ mutants and the parental cell line were subjected or not to total starvation for 2h in PBS, followed by sucrose gradient fractionation and polymer profile determination, where FP: free polysomes fraction, PP: pre-polysomes, LP: light polysomes and HP: heavy polysomes. (B) Accumulation of both RPL13a isoforms in light polysomes fraction was determined by western blotting after the starvation process and cycloheximide treatment. (C) Additionally, both RPL13a isoforms were found in light polysome fraction (LP), even after ribosome dissociation by puromycin treatment under normal and starvation conditions. Experiments were performed in duplicated and the polysome profiles correspond to the average of each sample.

## DISCUSSION

The relevance of post-transcriptional gene expression regulation in *Leishmania* parasites (17) associated to the typically duplicated genes encoding for RPs raised the hypothesis that such proteins might be subjected to different expression regulation mechanisms or even possess non-canonical functions. In our study, we observed a compensatory expression mechanism for the two pairs of RP paralogous genes, *RPS16* and *RPL13a* from *Leishmania major*, which is linked to an increase in either the rate of translation or protein stability of each paralogous protein, but without measurable effects on the levels of the corresponding transcripts. In humans, RPs are encoded by unique coding sequences, which regulates 10-20 copies of other genes, pseudogenes (32), whereas 59 genes encoding for RPs are found duplicated in the yeast genome (33). Gene duplication was considered as a redundancy in the genome, since, in general, carrying two identical copies of a given gene was thought not be advantageous (34). However, for highly expressed transcripts, as RP genes – the most expressed in the many cells (3) – such phenomenon can be favorable to supply the cell with high levels of RNA (34). Currently, gene duplication is understood as a recurrent and relevant process contributing to genome evolution (35). Cell survival in trypanosomatids is strongly conditioned to fast adaptation responses to hostile environments during the parasite life cyclev(19). Thus, agile mechanisms for regulation of gene expression are key for the survival of *Leishmania* parasites. However, a thorough understanding of these mechanisms is still missing.

*RPS16* and *RPL13a* genes were investigated at protein and transcript levels, aiming to uncover evidence and add information on the process of regulation of these genes. *RPS16* paralogs (*LmjF.26.0880* and *LmF.26.0890*) are localized *in tandem*, with identical CDSs and divergent *UTR*s (Fig1A and 1C). *RPL13a* genes (*LmjF.15.0200* and *LmjF.34.0860*), on the other hand, are found in different chromosomes, with divergent *UTR*s and non-identical CDSs (Fig1B and 1D). The divergent *UTR*s of paralogs may suggest different expression regulation for these genes since untranslated regions contain sites for the binding of RBP. 3’*UTR*s are strongly involved in the control of translational (29) and post-transcriptional (36) activities in eukaryotes. Also in prokaryotes, 3’*UTR*s are involved in gene expression regulation (37). All this evidence strongly suggests that the control of the expression for these duplicated genes may be directly related to divergences at the 3’*UTR*s.

When tagged at N-terminal (Fig2A), higher levels of one of the paralogs of RPS16 and RPL13a were always observed in both procyclic and metacyclic forms, compared to their respective copy (Fig2C). The abundance results observed for RPS16 isoforms corroborate those of RNA levels, previously observed by RT-qPCR (25). Due to the polycistronic nature of transcription in trypanosomatids, correlation between transcript and protein levels is loose. Also, in human cells, such correlation seems to be low, with only 33% of the genes showing some protein-mRNA levels correlation (38), being more commonly observed for stage specific genes (39). In this context, here we have demonstrated for the first time a positive correlation between RNA and protein levels for duplicated non-stage-specific mRNAs in *L. major* promastigotes.

Duplicated RPs might be associated with organism heterogeneity, and stage-specific RP isoforms were observed in *Arabidopsis thaliana*, with higher RPS5A level found in rapidly dividing cells during the early embryonic development, compared to its isoform, RPS5B, which is preferentially expressed in differentiating cells (40). In the same organism, RPL16B was another RP present in higher amounts in dividing cells, while the RPL16A expression shows a tissue-specific association (41). For *Leishmania* parasites, differentially expressed genes corroborate their agile adaptation to the different host environments during the life cycle. Interestingly, one of each paralogous RP genes displays transcript and protein levels characteristically at higher levels than their corresponding paralog, and adjustment of the protein amounts to native levels consistently happens to compensate for the absence of the other paralog. We also show that the paralog compensation mechanism for both RPS16 and RPL13a occurs at the protein level when their paralogous copy was deleted (Fig3B), similar to that observed for RPs from *Saccharomyces cerevisiae* (42). Such mechanism does not involve the control of the transcript steady-state (Fig3C). Thus, we may speculate that this compensatory expression mechanism happens because both isoforms play the canonical RP roles and that there is a fine-tuning of each protein level. As previously mentioned, the lack of regulators at individual genes delegates the control of gene expression in trypanosomatids to post-transcriptional processes, which includes mRNA steady-state, translation rate and protein stability or post-translational modifications (43). Untranslated regions of mRNAs play a central role on these processes as they contain sites for the binding of regulatory proteins, the RNA binding proteins (RBPs) (44,45). Herein, 11 and 16 proteins were identified as binding to the 3’*UTR*s of both *RPS16* and *RPL13a* isoforms, respectively (Fig4B). After gene ontology analysis, proteins involved in central aspects of translation, peptide and ribosome metabolism were identified in both 3’*UTR*s of *RPL13a* genes. For *RPS16*, most relevant pathways were related to the proteins involved in ribosomal biogenesis and metabolism (Fig4C). Thus, identified proteins in the *in vitro* pulldown assay reinforce the hypothesis that the compensatory mechanism for the duplicated transcripts involves the translation machinery. Comparing the pool of proteins, we observed that only 6 of them are common to all four RP 3’*UTR* sequences analyzed (Fig4C), and they might be the core RBPome for RP transcripts. Interestingly, two proteins were detected binding exclusively to the 3’*UTR*s of the isoforms present at lower levels, *RPL13_34* and *RPS16_90*, and eight proteins were only bound to the more abundant isoforms, *RPL13_15* and *RPS16_80*. The last two sets of proteins might be important for the specificities of control for the lower and higher RP expressors. In combination with these results, and aiming to find shared and specific *cis*-elements which might be binding sites for RBPs, we examined the 3’*UTR* sequences of the RP proteins and some conserved elements were found (FigSF4), deserving further experimental evaluation as putative functional binding sites implicated in the differential control of gene isoforms, as has been previously done by us (46).

To search for any possibility of non-conventional functions for the RPs here studied, we checked the nutritional stress response of mutant parasites, an essential process for the parasite adaptation and development in the in insect digestive tract (47). After 4h of starvation and 24h of recovery, MTT assays indicated higher cell viability for all the knockout parasite lines (Fig4A). Mitochondrial activity increment could be a result of long-term suffering signaling, and parasites are stressed even after 24 hours of recovery. This mitochondrial activity boost reveals a different nutritional stress resistance for the transfectants compared to the parental cell line. Nevertheless, we cannot disregard the limitations of MTT assay, as metabolic activity may be modified by external factors directly affecting the formazan conversion in culture (48). Although it cannot be ruled out at this point, we could not identify any non-canonical role for the RP isoforms investigated, but specific phenotypical alterations have been reported and associated with the absence of one copy of a duplicated RP, *uL6A* or *uL6B*, in *Saccharomyces cerevisiae*, resulting in variations in general protein synthesis (42). Furthermore, *RPL23AA* depletion was responsible to promote cell growth arrest with anomalies in *Arabidopsis thaliana*, even when the isoform *RPL23AB* is kept (49).

During starvation, *Leishmania* parasites usually store their mRNAs and some ribosomal constituents in cytoplasmic granules to protect them from degradation (50). The immunofluorescence images suggest that no changes on RPs distribution in the cell occurs under starvation (Fig5C), although a discrete, but detectable, clustering of these RPs suggests they are accumulating in some cytoplasmic aggregates (Fig5D). When starved, the presence of both RPL13a isoforms in light polysome fractions (Fig6B) indicates that these proteins are not exclusively associated to the ribosome subunits, even after total polysome dissociation by puromycin treatment (Fig6C). Such finding suggests that RPL13a isoforms may interact with mRNA molecules and accumulate with them as heavy complexes during nutritional stress. This idea is also supported by polysomic profile alteration for both *RPL13a* knockout parasites compared to the parental line (Fig6A). This finding could suggest that the depletion of either *RPL13a* isoforms results in a change in the translation machinery or in its stability, as based on the decrease of ribosomal subunits abundance. This is an indicative of canonical behavior for the *Leishmania* RPL13a since a similar pattern of polysomic profile distribution was observed for the *Trypanosoma brucei* RPL26, which plays a canonical function (28). However, in humans, RPL13a has been characterized as dispensable for ribosomal functionality, but essential for the mRNA methylation, a well-known non-canonical function for RP proteins (51). Other non-canonical functions have been also reported for the human RPL13a, playing different roles outside of the ribosome complex (52). These functions include the mRNA translation inhibition after RPL13a phosphorylation and its release from the ribosomal subunit (13) and as GAIT constituent, a protein-RNA complex which drives the selective transcription control for a group of related genes (53). Despite the lack of evidence of non-canonical functions for *L. major* RPL13a in our report, such hypothesis cannot be discarded since many reports have shown alternative roles for its orthologue in other organisms. Of note, differently from the Phenylalanine (Phe) and Glycine (Gly) present in L13a_15, their substitutes in the L13a_34 isoform, Cysteine (Cys) and Serine (Ser), are targets of a large variety of post-translational modifications (PTMs). As for serine, various PTMs such as phosphorylation, sulfation, and various sugar chain modifications may occur. Additionally, the nucleophilicity and redox-sensitivity characteristic of cysteine residues results in a variety of PTMs, including oxidation, nitrosylation, glutathionylation, prenylation, palmitoylation, and Michael adducts with lipid-derived electrophiles, or more rarely methylation and phosphorylation. Contrarily, Phe suffers no PTMs and Glycine is only subjected to N-Myristoylation or N-acetylation (at the N-terminus). These PTMs may regulate the activity, localization, and stability of a diversity of proteins (54). Nevertheless, many of these modifications are labile and dynamic, rendering it difficult to detect within a complex proteome (55). Therefore, a technical drawback or the conditions used to evaluate PTMs and moonlight activities for each of the L13a isoforms may explain our results. Yet, we must still consider this possibility given the characteristics of the amino acids substitutions.

Here, we have provided insights into the regulation of gene expression for duplicated genes in *Leishmania* parasites, which substantially indicates control at post-translational steps. The conserved *cis-*elements at 3’*UTR*s of these transcripts might be central to the compensatory mechanism observed at the protein level. RPL13a isoforms seem to play their canonical function as constituents in heavy complexes in the cell, critically remarkable under nutritional stress. However, a non-canonical role still requires deeper investigation to verify the potential of their orthologues in other organisms.

## ACKNOWLEDGEMENTS

We would like to thank Viviane Ambrosio for the technical support in the laboratory, the technical assistance at the microscopy facility (Ribeirão Preto Medical School), the staff of the Proteomics Platform of the CHU de Québec Research Centre, Québec, Canada and TriTrypDB.

